# Nectar non-protein amino acids (NPAAs) do not change nectar palatability but enhance learning and memory in honey bees

**DOI:** 10.1101/2020.12.04.407734

**Authors:** Daniele Carlesso, Stefania Smargiassi, Elisa Pasquini, Giacomo Bertelli, David Baracchi

**Author notes:** **Correspondence to** David Baracchi.

## Abstract

Floral nectar is a pivotal element of the intimate relationship between plants and pollinators and its chemical composition is likely to have been shaped by strong selective pressures. Nectars are composed of a plethora of nutritionally valuable compounds but also hundreds of secondary metabolites (SMs) whose ecological role is still not completely understood. Here we performed a set of behavioural experiments to study whether five ubiquitous nectar non-protein amino acids (NPAAs: β-alanine, GABA, citrulline, ornithine and taurine) interact with gustation, feeding preference, and learning and memory in the pollinator *Apis mellifera*. We showed that harnessed foragers were unable to discriminate NPAAs from water when only accessing antennal chemo-tactile information and that freely moving bees did not exhibit innate feeding preferences for NPAA-laced sucrose solutions. Also, dietary consumption of NPAAs did not alter food consumption or longevity in caged bees over 10 days. Taken together our data suggest that ecologically relevant concentrations of NPAAs did not alter nectar palatability to bees. Olfactory conditioning assays showed that honey bees were more likely to learn a scent when it signalled a sucrose reward containing either β-alanine or GABA, and that GABA also enhanced specific memory retention. Conversely, when ingested two hours prior to conditioning, GABA, β-alanine, and taurine weakened bees’ acquisition performances but not specific memory retention, which was enhanced in the case of β-alanine and taurine. Neither citrulline nor ornithine affected learning and memory. Our study suggests that NPAAs in nectars may represent a cooperative strategy adopted by plants to attract beneficial pollinators, while simultaneously enhancing pollen transfer among conspecific flowers. Future work should validate these results in more ecological scenarios and extend the study to as many nectar SMs as possible, alone and in combination, as well as to other species of pollinators.

## Introduction

Many plants depend on animal pollination for reproduction and they evolved flowers of a multitude of shapes, colours and fragrances that dispense sugary nectars to attract pollinators. In turn, pollinators feed on nectar and sustain plant reproduction by vectoring pollen among conspecific flowers [1, 2]. This classical ecological view of cooperative plant-pollinator interaction has been recently challenged by evidence of bilateral manipulative and cheating strategies [3, 4]. The emergence of these strategies has been promoted by conflicting interests: plants aim at maximizing their sexual reproduction while minimizing the costly production of nectar, whereas pollinators aim at maximizing food collection less concerned about pollen transfer. Floral nectar is a pivotal element of such an ambiguous interaction, and its chemical composition is likely to have been shaped by such strong selective pressures [5]. As a result, floral nectars are far from being a simple sugary reward. They are composed of a plethora of nutritionally valuable compounds, as well as hundreds of secondary metabolites (SMs), whose ecological role is still not completely understood [6-8].

Traditionally, the pleiotropy hypothesis has suggested that SMs, which include alkaloids, terpenes, phenolics and non-protein amino acids (NPAAs), are present in nectar only due to passive leakage from the plant’s phloem during transport to the surrounding tissues [2, 9, 10]. Recent research has suggested that nectar SMs may play a key role in plant-pollinator interactions by enhancing the quality and the quantity of pollination services received by plants [11-14]. For instance, it has been demonstrated that nectar alkaloids and diterpenes enhanced plant fitness by increasing pollen transfer among conspecific flowers, repelling nectar thieves and retaining specialist pollinators [12, 15-17]. This evidence, together with the fact that the composition of nectar precursors supplied to the nectaries is modified by selective secretion [18], progressively casted doubts on the pleiotropy hypothesis. The concentration of SMs in nectars is orders of magnitude lower than in other plant tissues, where their primary role is to deter herbivorous species from feeding [2, 9, 10]. Also, their impact on pollinators’ health and cognition is both species- and dose-dependent [19-24]. Despite the relevance that nectar SMs appear to have in plant-pollinator interactions, research on their ecological function is only limited to few alkaloids, a phenolic and a glycoside [15, 19-21, 25-27] whereas the role of other common SMs has been mostly overlooked.

Non-protein amino acids (NPAAs) are almost ubiquitous in flowering plants and represent the most abundant and frequent class of nectar SMs [7, 11]. As other SMs, NPAAs are not involved in primary metabolism, may possess antimicrobial properties and are toxic to animals when ingested in high doses [28, 29]. Although their distribution seems to be greatly variable across different plant taxa, γ-amino butyric acid (GABA) and β-alanine are certainly the most represented NPAAs in nectars [5, 11, 28]. A few other NPAAs, such as taurine, citrulline and ornithine, have also been often reported [11, 28]. Interestingly, NPAAs are most frequently found in hymenopteran-pollinated plant species and bees are the primary pollinators of those species that are abundant in GABA and ornithine [5, 11]. Taurine, GABA and β-alanine are also important neuromodulators in insects’ brain and are involved in muscle performance [30, 31]. GABA is the most abundant inhibitory neurotransmitter in the insect brain [32], and its function is essential for olfactory processing and learning [33, 34]. GABA receptors in insects are also located peripherally at the neuromuscular junctions [35], a fact that might explain why feeding on GABA-rich solutions affected general activity in both social and solitary bee species [36, 37]. Taken together, these data suggest that nectar NPAAs might have great relevance in regulating the relationship between plants and pollinators. However, since their discovery in nectars in the 1970s [7], no clear hypothesis has been put forward for their ecological role and their possible interactions with pollinators (but see [11] for a comprehensive review).

To fill this important lacuna, we performed a set of behavioural experiments to study whether five common nectar NPAAs interacted with gustation, feeding preferences, and learning in an insect pollinator. In our study, we tested GABA, β-alanine, taurine, citrulline and ornithine as they are the most common nectar NPAAs. We focused on the honey bee *Apis mellifera*, as it is a paradigmatic and valuable pollinator which offers the advantage of being a model system for studying behaviour and cognition [38, 39]. Bees display astonishing cognitive abilities and behavioural plasticity, which allow them to successfully navigate in complex environments, rapidly learn the cues associated with profitable food sources and communicate with nestmates [40, 41]. The learning abilities of bees can be studied in the laboratory through well-established standard procedures, such as the classical conditioning of the proboscis extension reflex (PER) [42]. In this protocol, a tethered bee is repeatedly presented with a neutral stimulus (CS, e.g. an odorant) paired with a sucrose reinforcement (US), which innately evokes the PER. This way, the animal will learn the association between the stimuli and, at the end of the training, the conditioned stimulus (CS) will elicit the PER response by itself [42, 43]. This protocol has been widely used in neurobiological and toxicological studies, and thus represents a standard method to explore how external substances influence bees’ cognition. In a first experiment, we used a chemo-tactile conditioning of PER to investigate whether bees could detect ecologically relevant concentrations of nectar NPAAs when using only their antennae. As bees may assess nectar quality through chemosensilla located on their mouthparts [44, 45], we measured whether freely moving bees showed innate feeding preferences for NPAA-laced sucrose solutions over a two-minute binary choice assay. Nectar secondary metabolites may also induce post-ingestive malaise or phagostimulation over longer periods of times [8, 44, 46]. We therefore investigated whether dietary consumption of NPAAs affected food consumption and longevity in caged bees over 10 days. Lastly, we used a series of olfactory PER conditioning assays to evaluate whether NPAAs affected associative learning and memory in bees, either when NPAAs were used as a reward during conditioning or when NPAAs were ingested prior to training.

## Results

### Exp 1 – Chemo-tactile conditioning of the proboscis extension response (PER)

Bee foragers may assess the quality of floral nectars through chemo-sensilla located on their antennae [47]. In this first experiment, we asked whether nectar-relevant concentrations of five common NPAAs (i.e. GABA, β-alanine, taurine, citrulline and ornithine) can be detected by bees through their antennae. To this aim, we used a chemo-tactile differential conditioning of PER protocol [48] in which different groups of bees were trained to discriminate one of the five NPAAs from water. Briefly, tethered bees experienced five pairings of a neutral stimulus (either NPAA-laced water or water) (CS+) with a 30% sucrose solution reinforcement (US) and five pairings (either water or NPAA-laced water) (CS-) with a saturated NaCl solution (US) used as punishment. If an association is formed, the conditioned stimulus (CS+) should elicit the PER by itself [42, 43]. Note that learning could occur only if bees detected the NPAA in the solution through their antennae. CSs were presented to the bees’ antennae via a small piece of filter paper soaked in the appropriate solution and were presented in a pseudorandom order according to their contingency (i.e. rewarded *vs.* punished). The results showed that bees increased their response to both the rewarded (CS+) and the punished (CS-) stimuli over the ten conditioning trials (GLMM, *trial*: GABA: n = 76, *χ*^*2*^ = 65.75, df = 1, p < 0.0001; β-alanine: n = 81, *χ*^*2*^ = 98.15, df = 1, p < 0.0001; ornithine: n = 72, *χ*^*2*^ = 39.23, df = 1, p < 0.0001; taurine: n = 69, *χ*^*2*^ = 59.24, df = 1, p < 0.0001; citrulline: n = 79, *χ*^*2*^ = 60.70, df = 1, p < 0.0001, Fig. 1). In all cases, the responses to the CSs did not differ over the course of the training, suggesting that bees were not able to discriminate any dissolved NPAAs from water with their antennae (GLMM, *CS*: GABA: *χ*^*2*^ = 3.01, df = 1, p = 0.08; β-alanine: *χ*^*2*^ = 2.68, df = 1, p = 0.10; ornithine: *χ*^*2*^ = 3.31, df = 1, p = 0.07; taurine: *χ*^*2*^ = 0.20, df = 1, p = 0.65; citrulline: *χ*^*2*^ = 1.89, df = 1, p = 0.17, Fig. 1).

**Figure 1:**
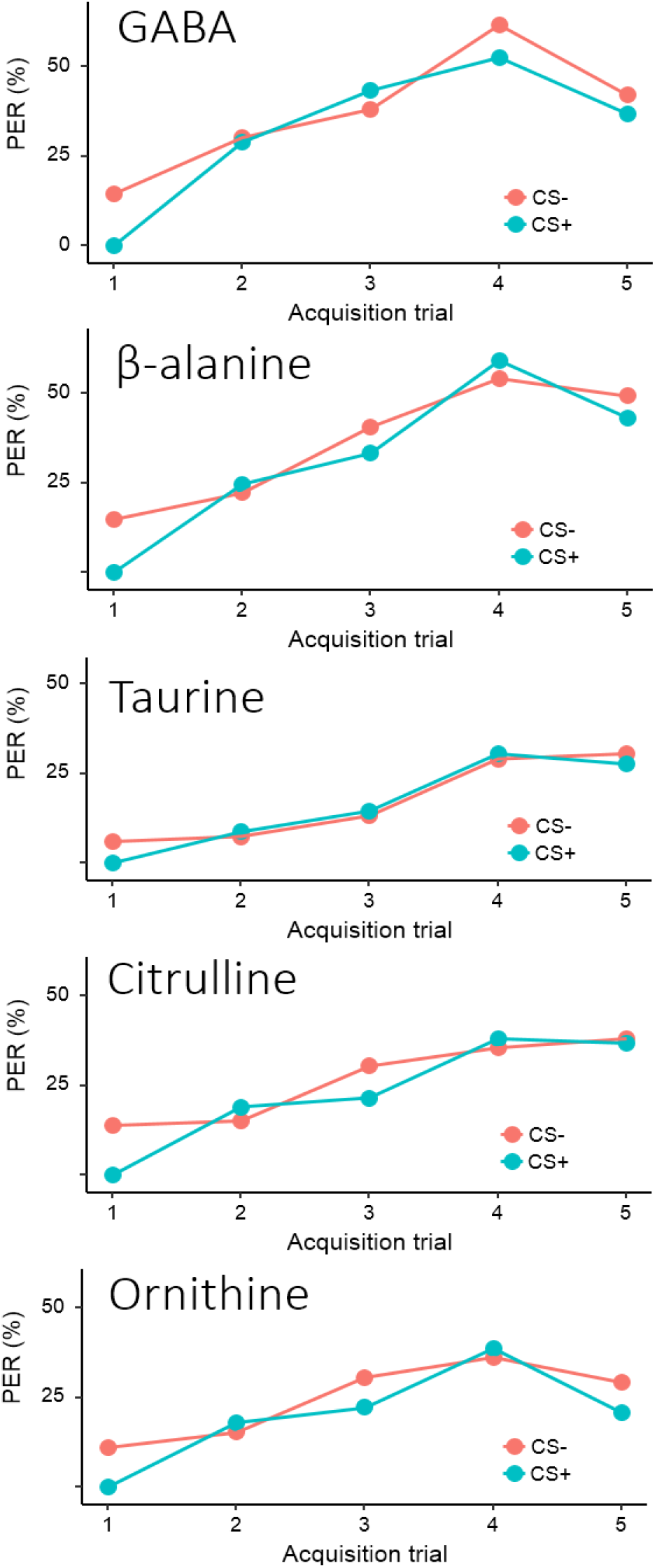
Honey bees are not able to detect NPAAs when using only their antennae. Proportion of bees showing a conditioned PER during the chemo-tactile conditioning assay. Blue lines represent PER to the reinforced stimulus (CS+). Pink lines represent PER to the punished stimulus (CS-). NPAA-laced water solutions and pure water were used as CS+ and CS-. No significant differences were found in the proportion of bees showing PER to the CS+ and the CS- for any of the NPAAs (GLMM, GABA: p = 0.07; β-alanine: p = 0.10; taurine: p = 0.65; citrulline: p = 0.17; ornithine: p = 0.07).

### Exp 2 – Taste aversion/preference assay

Besides using their antennae, bees may detect nectar constituents through the chemo-sensilla located on their mouthparts [44, 45]. We thus further investigated the gustatory responses of bees to nectar NPAAs using a binary choice protocol adapted from Ma and colleagues [49]. After one hour resting, freely moving bees confined into custom-modified plastic tubes were presented with two microcapillaries, one containing 100 μl of 30% sucrose solution and the other containing 100 μl of NPAA-laced 30% sucrose solution. We recorded the feeding behaviour of each bee for two minutes after the first approach to one of the solutions. NPAAs were always tested against plain sucrose and independent groups of bees were used for each NPAA (GABA, n = 56; β-alanine, n = 46; taurine, n = 55; citrulline, n = 57; ornithine, n = 58). A preference index for each NPAA was calculated by subtracting the residual volume of the liquid inside the two capillaries at the end of test. The comparison of the preference index against the hypothetical value of 0, which indicates a lack of preference, showed that bees did not exhibit a feeding preference nor avoidance for any of the nectar NPAAs (One-Sample Wilcoxon test, GABA: V = 796, p = 0.99; β-alanine: V = 512, p = 0.76; taurine: V = 855, p = 0.48; citrulline: V = 925, p = 0.44; ornithine: V = 911, p = 0.67, Fig. 2).

**Figure 2:**
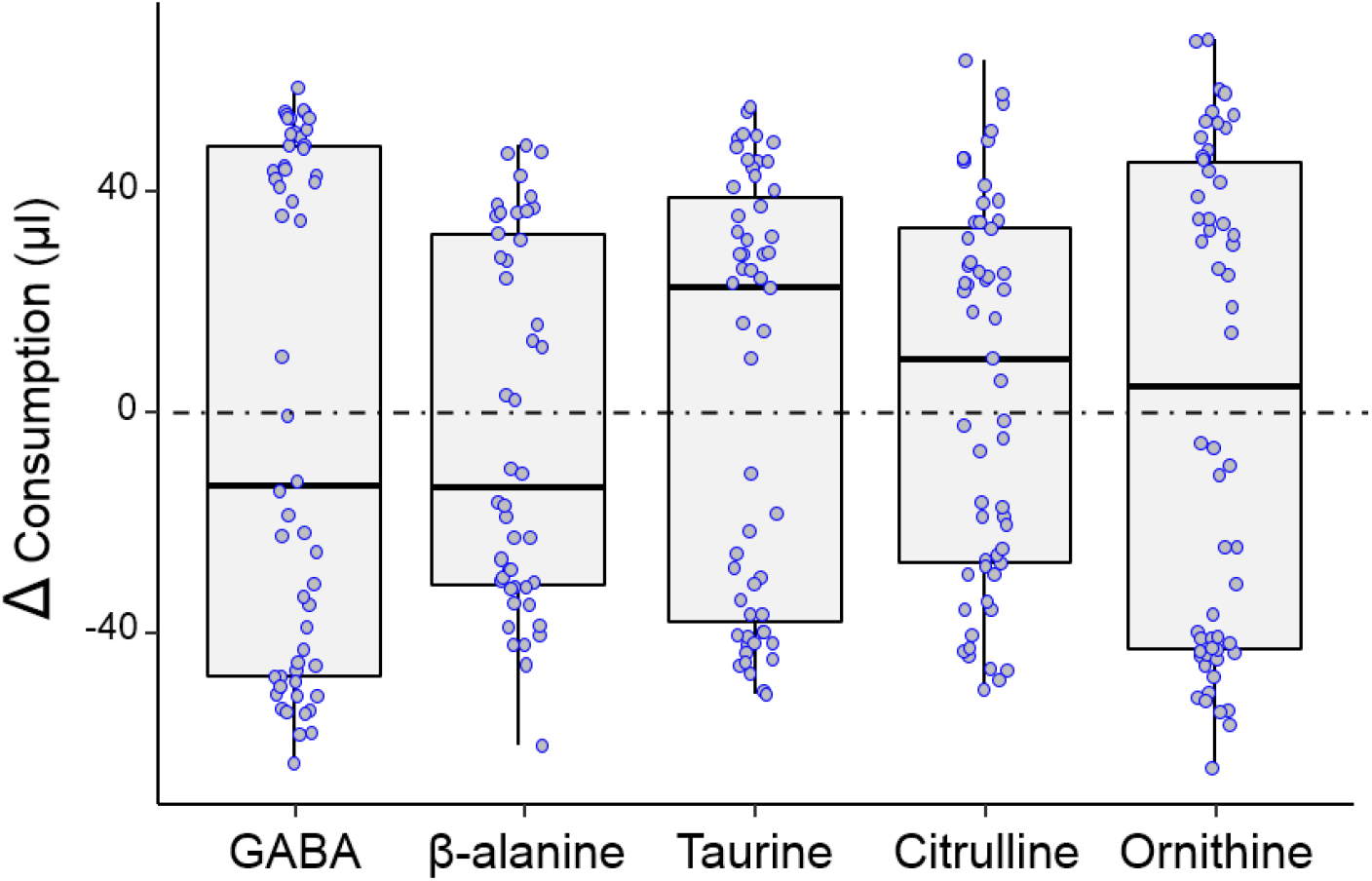
NPAAs are not preferred or avoided by honey bees in the feeding assay. Median, quartiles, minimum and maximum consumption indexes observed for each of the NPAAs in the feeding preference test. A higher Δ value indicates a preference for the NPAA-laced sucrose solution. A lower Δ value indicates a preference for plain sucrose solution. Dots represent individual bees. Bees did not exhibit any clear preference or avoidance for any of the NPAAs (One-Sample Wilcoxon test, GABA: p = 0.99; β-alanine: p = 0.76; taurine: p = 0.48; citrulline: p = 0.44; ornithine: p = 0.67).

### Exp 3 – Influence of NPAAs on feeding and mortality

In the previous experiment, we showed that bees did not have a reflexive (pre-ingestive) taste aversion or preference towards NPAAs. However, food aversion/preference may eventually arise from malaise or phagostimulation caused by SM ingestion [28, 44, 46]. To test this hypothesis, we housed a total of 900 bees in groups of 10 individuals each in plastic cages equipped with two food dispensers (syringes) [50]. For each NPAA, six replicates for three experimental conditions were set up: 1) two syringes providing plain sucrose solution (S-S); 2) two syringes providing NPAA-laced sucrose solution (NPAA-NPAA); 3) one syringe providing plain sucrose solution and another one providing NPAA-laced sucrose solution (S-NPAA). The experiment lasted until all bees in the cages were dead. The results showed that bees kept under the S-S and the NPAA-NPAA regimes did not differ in *per capita* food consumption at 24 hours from the beginning of the experiment (Mann-Whitney U test, GABA, p = 0.75; β-alanine, p = 0.59; taurine, p = 0.31; citrulline, p = 0.70; ornithine, p = 0.59). However, taurine decreased the overall amount of food consumed over 10 days (Mann-Whitney U test, p = 0.009). All the other NPAAs had no effect on the *per capita* total amount of food consumed (GABA, p = 0.81; β-alanine, p = 0.94; citrulline, p = 0.70; ornithine, p = 0.94). Bees kept under the S-NPAA regime could choose between pure sucrose and sucrose laced with one of the five NPAAs. Yet, bees showed no preference/avoidance for the NPAAs at 24 hours (Mann-Whitney U test, GABA, p = 0.75; β-alanine, p = 0.39; taurine, p = 0.91; citrulline, p = 0.75; ornithine, p = 0.89) nor at the end of the experiment (Mann-Whitney U test, GABA, p = 0.35; β-alanine, p = 0.46; taurine, p = 0.14; citrulline, p = 0.50; ornithine, p = 0.29). A statistical evaluation of bees’ survival during the experiment revealed that NPAAs were not a significant predictor of mortality in any feeding regime (Log-rank Mantel Cox test, GABA, *χ*^*2*^ = 0.77, df = 2, p = 0.68; β-alanine, *χ*^*2*^ = 1.32, df = 2, p = 0.52; taurine, *χ*^*2*^ = 0.78, df = 2, p = 0.68; citrulline, *χ*^*2*^ = 3.19, df = 2, p = 0.20; ornithine, *χ*^*2*^ = 2.80, df = 2, p = 0.25, Fig. 3).

**Figure 3:**
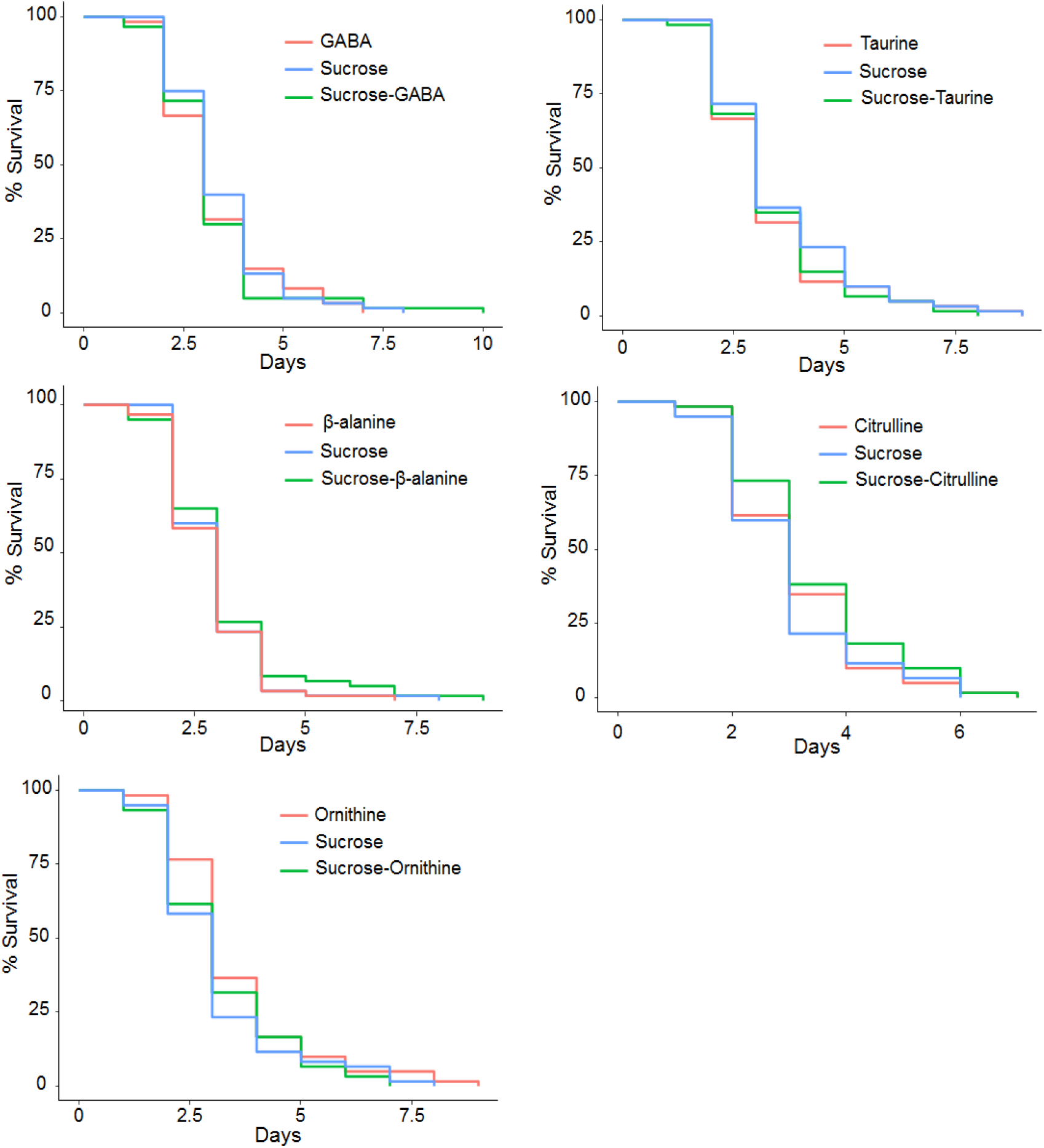
NPAAs did not affect caged honey bee survival. Cumulative survival of bees kept in caged conditions under three different feeding regimes for a period of 10 days: Sucrose only (S-S); NPAA-laced solution only (NPAA-NPAA); Sucrose and NPAA-laced solution (S-NPAA). None of the NPAAs was a significant predictor of mortality in any feeding regime (Log-rank Mantel Cox test, GABA: p = 0.68; β-alanine: p = 0.52; taurine: p = 0.68; citrulline: p = 0.20; ornithine: p = 0.25).

### Exp 4 – Contextual absolute olfactory learning

Nectar metabolites may enhance or inhibit bees’ ability to learn floral cues by directly acting on the nervous system, regardless of their ability to trigger gustatory and olfactory receptor neurons of bees’ antennae and mouthparts [14, 31]. Thus, we investigated whether the presence of NPAAs in sucrose rewards altered honey bees’ learning and memory performance during an olfactory absolute conditioning task [42, 43]. Briefly, for each of the five NPAAs, tethered bees were presented with four pairings of a neutral odorant (CS, either 1-Hexanol or Nonanal) with a reinforcement delivered to the antennae and mouthparts (US, either 30% sucrose solution (*control paired group*) or NPAA-laced 30% sucrose solution (*experimental paired group*)). In addition, for each of the five NPAAs, we trained two *explicitly unpaired* groups of bees to control for true associative learning [42]. In this case, bees experienced four presentations of the CS and four presentations of either the NPAA-laced or plain sucrose solution but in absence of pairing, so that they could not form any association between the stimuli [43]. All bees in the *paired* groups increased their responses to the conditioned odorant over the four training trials (GLMM, *trial*: GABA, *χ*^*2*^ = 57.9, df = 1, p < 0.0001; β-alanine, *χ*^*2*^ = 52.3, df = 1, p < 0.0001; taurine, *χ*^*2*^ = 42.1, df = 1, p < 0.0001; citrulline, *χ*^*2*^ = 53.3, df = 1, p < 0.0001; ornithine, *χ*^*2*^ = 52.1, df = 1, p < 0.0001, Fig. 4). GABA and β-alanine significantly enhanced acquisition performances in bees (GLMM, *treat*: GABA, *χ*^*2*^ = 4.87, df = 1, p = 0.027; β-alanine, *χ*^*2*^ = 6.86, df = 1, p = 0.009, Fig. 4). Conversely, no effect on learning was observed for taurine, citrulline and ornithine (GLMM, *treat:* taurine, *χ*^*2*^ = 0.003, df = 1, p = 0.95; citrulline, *χ*^*2*^ = 0.84, df = 1, p = 0.36; ornithine, *χ*^*2*^ = 0.58, df = 1, p = 0.45). Bees that received GABA or β-alanine as reward also had significantly higher acquisition scores (ACQS, calculated as the number of PER exhibited over the 4 trials) than controls (Mann-Whitney U test, *ACQS:* GABA, W = 922, p = 0.041; β-alanine, W = 1495, p = 0.006). No difference was observed for any of the other NPAAs (Mann-Whitney U test, *ACQS*: taurine, W = 707, p = 0.87; citrulline: W = 1007, p = 0.34; ornithine: W = 799, p = 0.39). Our analysis showed that the conditioned stimulus (CS, either Nonanal or 1-Hexanol) significantly affected responses of both experimental and control bees in both the *paired* and *unpaired* GABA groups. In particular, bees in the *paired* groups showed a significantly higher number of responses toward 1-Hexanol than to Nonanal (GLMM, *CS*: χ^2^ = 27.22, df = 1, p < 0.0001), whereas the contrary was true in the *unpaired* groups (GLMM, *CS*: *χ*^*2*^ = 4.4, df = 1, p = 0.036). For all *unpaired* groups, we observed no significant increase in responses during conditioning apart from β-alanine (GLMM, *trial*: GABA, *χ*^*2*^ = 0.15, df = 1, p = 0.70; β-alanine, *χ*^*2*^ = 4.32, df = 1, p = 0.04; taurine, *χ*^*2*^ = 1.26, df = 1, p = 0.26; citrulline, *χ*^*2*^ = 0.19, df = 1, p = 0.66; ornithine, *χ*^*2*^ = 0.59, df = 1, p = 0.44, Fig. 4), nor an effect of the treatment (GLMM, *treat:* GABA, *χ*^*2*^ = 0.04, df = 1, p = 0.84; β-alanine, *χ*^*2*^ = 0.13, df = 1, p = 0.71; taurine, *χ*^*2*^ = 0.01, df = 1, p = 0.93; citrulline, *χ*^*2*^ = 0.0, df = 1, p = 0.99; ornithine, *χ*^*2*^ = 0.001, df = 1, p = 0.98, Fig. 4). Accordingly, bees in the *unpaired* groups did not differ in acquisition scores for any of the NPAAs (Mann-Whitney U test, *ACQS:* GABA, W = 771, p = 0.92; β-alanine, W = 818, p = 0.39; taurine, W = 740, p = 0.15; citrulline, W = 861, p = 0.34; ornithine, W = 722, p = 0.1). These results thus confirmed the occurrence of a true associative learning phenomenon in the *paired* groups.

**Figure 4:**
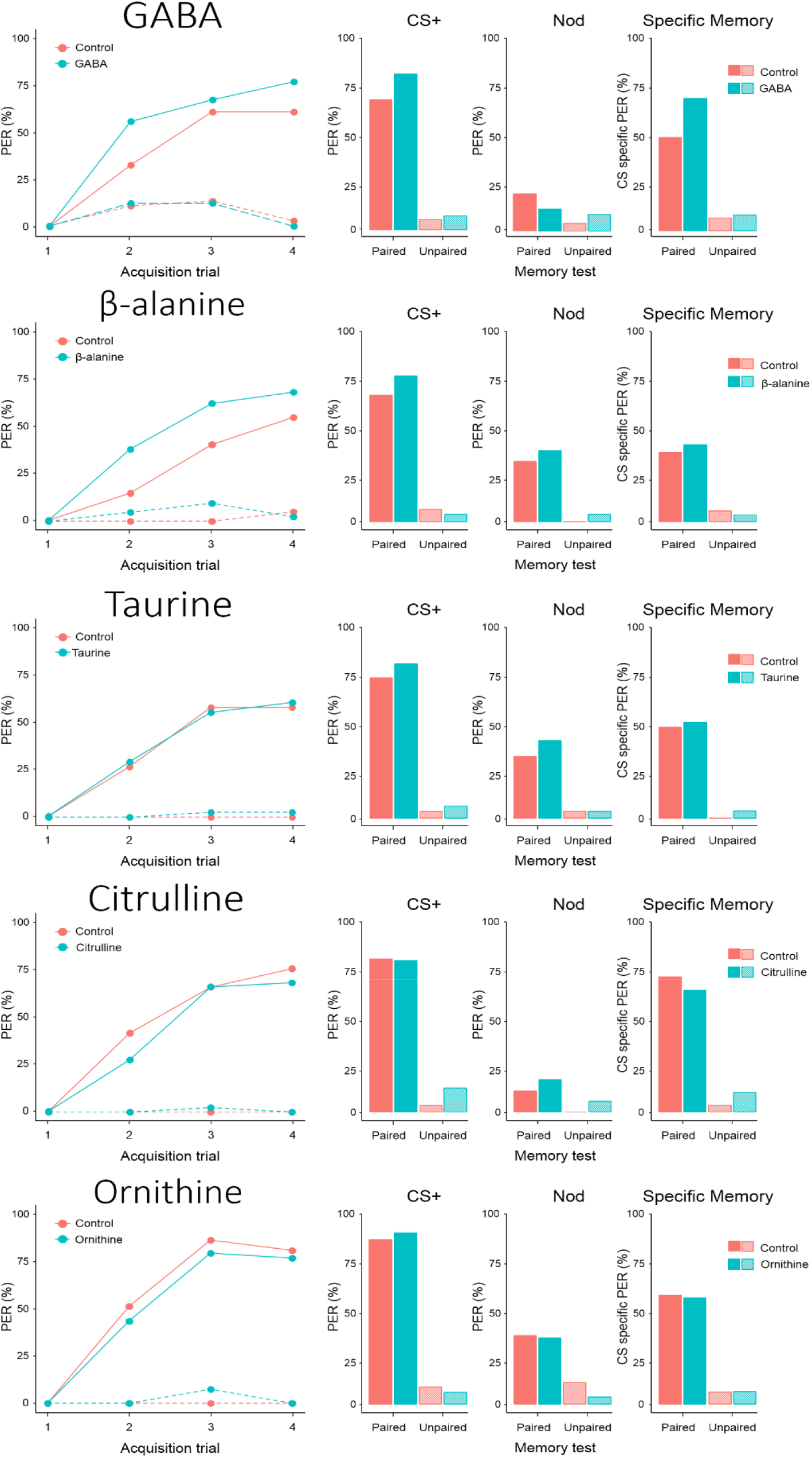
Effects of NPAAs dissolved in US on appetitive olfactory learning and memory retention in harnessed bees. **Acquisition trials:** Percentages of PER showed by experimental (blue lines) and control (red lines) bees in the paired (solid lines) and unpaired (dotted lines) groups during the contextual conditioning experiment for each of the NPAAs. In the paired groups, GABA and β-alanine significantly enhanced the acquisition performances of bees (GLMM, GABA, p = 0.027; β-alanine, p = 0.009). None of the other NPAAs affected learning performances (all cases: p > 0.4). Unpaired groups did not learn the US-CS association (all cases: p > 0.7). **Memory test:** Proportions of PER showed by experimental (blue) and control (red) bees in the paired and unpaired groups during the memory test performed two hours after conditioning. GABA significantly enhanced the specific memory of bees for the trained odorant (*χ*^*2*^ test, p = 0.03). No difference has been found in the responses to the CS+, to the NOd or in CS-specific memory in all other cases (all cases: p > 0.1). Unpaired groups did not differ in the memory performances (all cases: p > 0.2).

Two hours after conditioning, bees were tested for memory retention. In this non-reinforced memory test, bees were presented once with the conditioned odorant (CS, either 1-Hexanol or Nonanal) and once with a novel odorant (NOd, either 1-Hexanol or Nonanal depending on CS) to investigate whether bees’ responses were CS-specific or generalized [43]. Experimental and control bees in all the *paired* groups did not differ in their responses to the conditioned odorant for any of the NPAAs (χ^2^ test, *CS:* GABA, *χ*^*2*^ = 2.3, p = 0.13; β-alanine, *χ*^*2*^ = 1.9, p = 0.16; taurine, *χ*^*2*^ = 0.68, p = 0.41; citrulline, *χ*^*2*^ = 0.01, p = 0.91; ornithine, *χ*^*2*^ = 0.19, p = 0.66, Fig. 4). Neither did responses to the novel odorant (NOd) differ between experimental and control bees (*χ*^*2*^ test, *NOd:* GABA, *χ*^*2*^ = 1.21, p = 0.27; β-alanine, *χ*^*2*^ = 0.47, p = 0.49; taurine, *χ*^*2*^ = 0.52, p = 0.47; citrulline, *χ*^*2*^ = 0.71, p = 0.40; ornithine, *χ*^*2*^ = 0.27, p = 0.87, Fig. 4). GABA significantly enhanced specific memory for the conditioned odorant (*χ*^*2*^ test, *specific memory*: *χ*^*2*^ = 4.63, p = 0.03, Fig. 4). No such difference was found for any of the other nectar NPAAs (*χ*^*2*^ test, *specific memory*: β-alanine, *χ*^*2*^ = 0.40, p = 0.53; taurine, *χ*^*2*^ = 0.053, p = 0.82; citrulline, *χ*^*2*^ = 0.48, p = 0.49; ornithine, *χ*^*2*^ = 0.001, p = 0.98, Fig. 4). As expected, in all the *unpaired* groups NPAAs did not alter bees’ responses to the conditioned odorant (*χ*^*2*^ test, *CS:* GABA, *χ*^*2*^ = 0.14, p = 0.71; β-alanine, *χ*^*2*^ = 0.37, p = 0.54; taurine, *χ*^*2*^ = 0.38, p = 0.55; citrulline, *χ*^*2*^ = 2.67, p = 0.10; ornithine, *χ*^*2*^ = 0.24, p = 0.62, Fig. 4) nor to the novel odorant (*χ*^*2*^ test, *NOd:* GABA, *χ*^*2*^ = 0.90, p = 0.34; β-alanine, *χ*^*2*^ = 0.99, p = 0.32; taurine, *χ*^*2*^ = 0.0003, p = 0.99; citrulline, *χ*^*2*^ = 1.95, p = 0.16; ornithine, *χ*^*2*^ = 2.00, p = 0.16, Fig. 4). Accordingly, we found no difference in the proportion of experimental and control bees showing CS-specific memory for any of the NPAAs (*χ*^*2*^ test, *specific memory*: GABA, *χ*^*2*^ = 0.14, p = 0.71; β-alanine, *χ*^*2*^ = 0.37, p = 0.54; taurine, *χ*^*2*^ = 1.04, p = 0.31; citrulline, *χ*^*2*^ = 1.77, p = 0.18; ornithine, *χ*^*2*^ = 0.001, p = 0.98, Fig. 4).

### Exp 5 – Post-feeding absolute olfactory learning

In the previous experiment, we demonstrated that GABA and β-alanine altered learning in bees when ingested during the conditioning phase. Here we explored the post-ingestive mechanisms of NPAAs on bees’ learning and memory performance. In this case, bees were fed 5 μL of either NPAA-laced 30% sucrose solution (*experimental groups*) or plain 30% sucrose solution (*control groups*) two hours before conditioning. Bees were then confronted with a conditioning procedure identical to that described above (Exp. 4), except that we always used plain sucrose as US. As before, for each NPAA we trained a *paired* and an *explicitly unpaired* groups to verify true associative learning [43] and an individual acquisition score (ACQS) was calculated for each bee. Bees in all the *paired* groups significantly increased their responses to the conditioned odorant over the four training trials (GLMM, *trial*: GABA, *χ*^*2*^ = 39.5, df = 1, p < 0.0001; β-alanine, *χ*^*2*^ = 35.35, df = 1, p < 0.0001; taurine, *χ*^*2*^ = 46.8, df = 1, p < 0.0001; citrulline, *χ*^*2*^ = 51.7, df = 1, p < 0.0001; ornithine, *χ*^*2*^ = 56.3, df = 1, p < 0.0001, Fig. 5). Bees pre-fed GABA and taurine had significantly worse acquisition performances than controls (GLMM, *treat*: GABA, *χ*^*2*^ = 5.79, df = 1, p = 0.016; taurine, *χ*^*2*^ = 3.91, df = 1, p = 0.048, Fig. 5). β-alanine had a similar almost significant effect (GLMM, *treat*: β-alanine, *χ*^*2*^ = 2.73, df = 1, p = 0.098). Neither ornithine nor citrulline had an effect on learning performances (GLMM, *trial:* citrulline, *χ*^*2*^ = 1.83, df = 1, p = 0.18; ornithine, *trial: χ*^*2*^ = 0.01, df = 1, p = 0.99, Fig. 5). Accordingly, ACQS were significantly lower in bees pre-fed GABA than controls (Mann-Whitney U test, W = 656, p = 0.035). A non-significant tendency in the same direction was found for both β-alanine (Mann-Whitney U test, W = 817, p = 0.077) and taurine (Mann-Whitney U test, W = 833, p = 0.057). No difference in ACQS values was found between experimental and control bees pre-fed citrulline (Mann-Whitney U test, W = 967, p = 0.13) or ornithine (Mann-Whitney u test, W = 807, p = 0.94). Bees belonging to the *unpaired* groups did not increase their responses over training except with β-alanine (GLMM, *trial:* GABA, *χ*^*2*^ = 3.56, df = 1, p = 0.06; β-alanine, *χ*^*2*^ = 4.87, df = 1, p = 0.03; taurine, *χ*^*2*^ = 0.38, df = 1, p = 0.54; citrulline, *χ*^*2*^ = 1.26, df = 1, p = 0.26; ornithine, *χ*^*2*^ = 2.87, df = 1, p = 0.1, Fig. 5). However, in all the *unpaired groups*, including β-alanine, pre-feeding did not alter the responses to the CS (GLMM, *treat:* GABA: *χ*^*2*^ = 0.04, df = 1, p = 0.84; β-alanine: *χ*^*2*^ = 1.88, df = 1, p = 0.17; taurine: *χ*^*2*^ = 0.0004, df = 1, p = 0.98; citrulline: *χ*^*2*^ = 0.06, df = 1, p = 0.81; ornithine: *χ*^*2*^ = 0.053, df = 1, p = 0.82, Fig. 5). Accordingly, experimental and control pre-fed bees did not differ in their ACQS (Mann-Whitney U test, *ACQS*: GABA: W = 529, p = 0.69; β-alanine: W = 559, p = 0.15; taurine: W = 666, p = 1; citrulline: W = 841, p = 0.98; ornithine: W = 665, p = 0.68). Overall, the results confirmed that true associative learning occurred in the *paired* groups.

**Figure 5:**
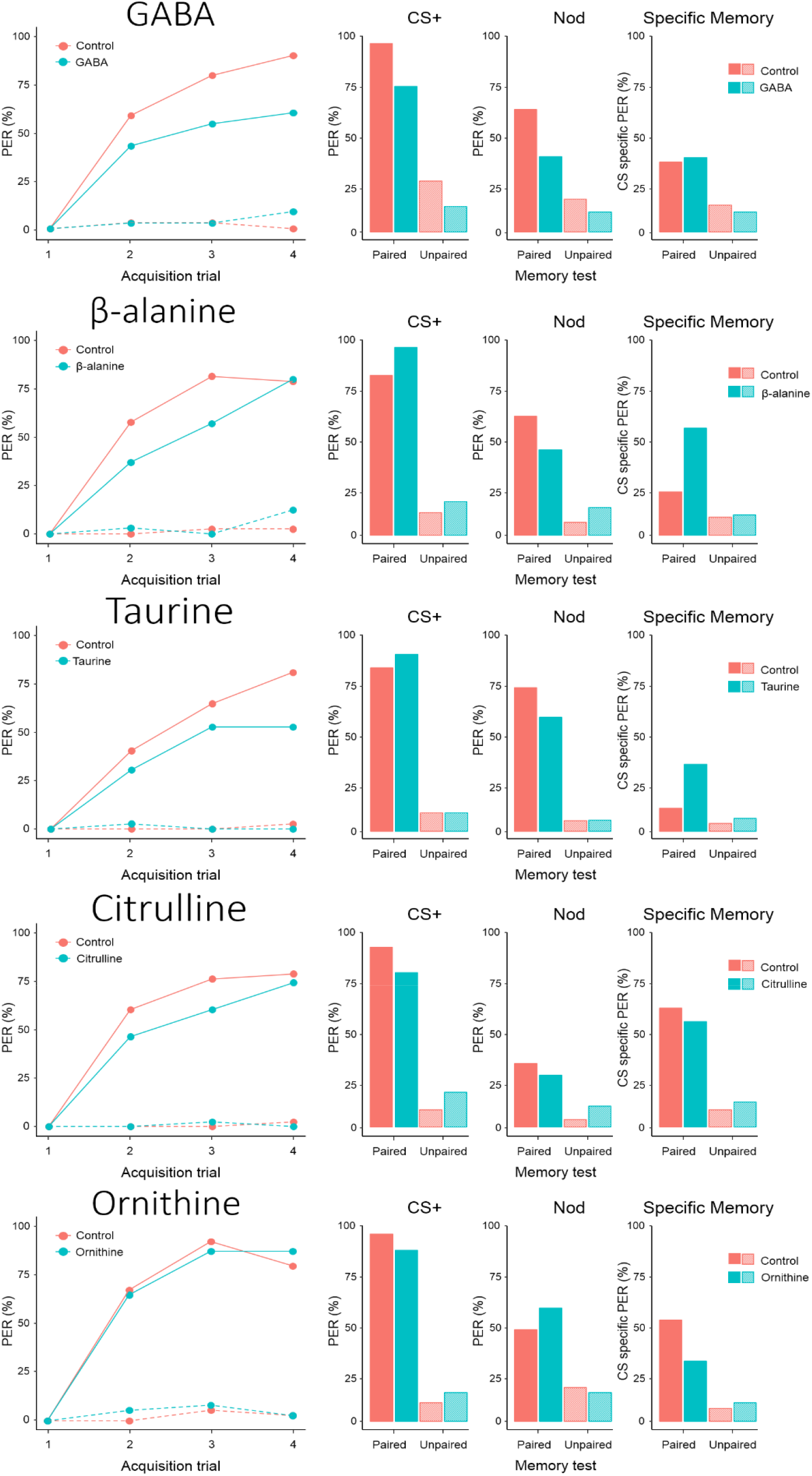
Effects of pre-feeding NPAAs on appetitive olfactory learning and memory retention in harnessed bees. **Acquisition trials:** Percentages of PER exhibited by experimental (blue lines) and control (red lines) bees in the paired (solid lines) and unpaired (dotted lines) groups during the post-feeding absolute olfactory learning experiment. Pre-ingestion of GABA and taurine impaired acquisition performance in honey bees (GLMM, GABA, p = 0.016; taurine, p = 0.048). β-alanine had a similar almost significant effect (p = 0.098). Ingestion of citrulline and ornithine did not affect associative learning (citrulline, p = 0.18; ornithine, p = 0.99). Unpaired groups did not learn the US-CS association (all cases: p > 0.17). **Memory test:** proportions of PER showed by experimental (blue) and control (red) bees in the paired and unpaired groups during the memory retention test performed two hours after the post-feeding conditioning assay. In the paired groups, bees pre-fed GABA exhibited a lower number of PER to the CS (*χ*^*2*^ test, p = 0.01) and to the novel odorant (p = 0.047). However, GABA did not affect bees’ specific memory (p = 0.83). β-alanine and taurine enhanced bees’ specific memory (β-alanine, p = 0.003; taurine, p = 0.02). In all other cases there were no significant differences in the response to the CS+, to the NOd and in the specific memory (all cases, p > 0.1). Unpaired groups did not differ in the memory performances (all cases: p > 0.2).

Two hours after conditioning, bees were tested for memory retention. As in the previous experiment, the CS and the novel odorant (NOd) were presented to the bees without reward. Bees pre-fed GABA showed significantly less appetitive responses to the CS (χ^2^ test, *χ*^*2*^ = 5.96, p = 0.01) and to the novel odorant (*χ*^*2*^ test, *χ*^*2*^ = 3.95, p = 0.047) than controls (Fig. 5). However, their CS-specific memory was not affected compared to control bees (*χ*^*2*^ test, *χ*^*2*^ = 0.49, p = 0.83, Fig. 5). Bees pre-fed β-alanine exhibited a significantly higher number of appetitive responses to the CS than controls (*χ*^*2*^ test, *χ*^*2*^ = 4.52, p = 0.03) but not to the novel odorant (*χ*^*2*^ test, *χ*^*2*^ = 2.28, p = 0.13, Fig. 5). Accordingly, a higher proportion of bees pre-fed β-alanine exhibited CS-specific memory (*χ*^*2*^ test, *χ*^*2*^ = 8.63, p = 0.003, Fig. 5). Taurine, citrulline and ornithine did not alter bees’ response to the CS (*χ*^*2*^ test, taurine: *χ*^*2*^ = 0.40, p = 0.53; citrulline: *χ*^*2*^ = 2.72, p = 0.1; ornithine: *χ*^*2*^ = 2.78, p = 0.09) or to the novel odorant (*χ*^*2*^ test, taurine: *χ*^*2*^ = 1.74, p = 0.19; citrulline: *χ*^*2*^ = 0.36, p = 0.55; ornithine: *χ*^*2*^ = 1.01, p = 0.31, Fig. 5). However, a significantly higher proportion of bees pre-fed taurine exhibited CS-specific memory during the test (*χ*^*2*^ test, *χ*^*2*^ = 5.41, p = 0.02). Citrulline and ornithine had no effect on memory retention (*χ*^*2*^ test, Citrulline: *χ*^*2*^ = 0.41, p = 0.52; Ornithine: *χ*^*2*^ = 3.71, p = 0.054). In the *unpaired* groups, no NPAA altered the responses to the CS (*χ*^*2*^ test, GABA: *χ*^*2*^ = 0.94, p = 0.16; β-alanine: *χ*^*2*^ = 0.53, p = 0.47; taurine: *χ*^*2*^ = 0.001, p = 0.97; citrulline: *χ*^*2*^ = 1.61, p = 0.20; ornithine: *χ*^*2*^ = 0.56, p = 0.45) or to the NOd (*χ*^*2*^ test, GABA: *χ*^*2*^ = 0.72, p = 0.40; β-alanine: *χ*^*2*^ = 1.33, p = 0.25; taurine: *χ*^*2*^ = 0.001, p = 0.98; citrulline: *χ*^*2*^ = 1.77, p = 0.18; ornithine: *χ*^*2*^ = 0.11, p = 0.74, Fig. 5). Accordingly, *unpaired* experimental and control bees did not differ in CS-specific memory for any of the NPAAs (*χ*^*2*^ test, GABA: *χ*^*2*^ = 0.24, p = 0.63; β-alanine: *χ*^*2*^ = 0.049, p = 0.83; taurine: *χ*^*2*^ = 0.38, p = 0.54; citrulline: *χ*^*2*^ = 0.45, p = 0.50; ornithine: *χ*^*2*^ = 0.21, p = 0.64, Fig. 5).

## Discussion

Here we report our results showing the impact of nectar-relevant concentrations of NPAAs on honey bee cognition. In our first behavioural assay, bee foragers were unable to detect dissolved NPAAs when only antennal chemo-tactile information was available. We also demonstrated that bees did not exhibit innate feeding preference between a plain sucrose solution and an equal solution laced with NPAAs in a two-minute assay. In line with a previous study [36], our 10-day-long toxicological assays showed that nectar NPAAs affected neither food consumption nor bees’ longevity. Taken together, these results strongly suggest that nectar NPAAs at concentrations within the natural range do not alter nectar palatability, at least for the pollinator *Apis mellifera*. Interestingly, we found that honey bees were more likely to learn a scent when it signalled a sucrose reward containing either β-alanine or GABA. GABA also enhanced specific memory retention. Conversely, when ingested two hours prior to conditioning, GABA, β-alanine, and taurine weakened bees’ acquisition performances but not the retention of the olfactory information, which actually became enhanced in the case of β-alanine and taurine.

Positive or negative effects on learning performances in bees have already been reported for several nectar SMs, including phenolics and alkaloids such as quercetin, naringenin, nicotine and caffeine [19, 20, 27, 51]. In our study honey bees were significantly better at learning scents signalling rewards containing either GABA or β-alanine in an olfactory PER classical conditioning assay. By contrast, the other tested NPAAs impacted neither the acquisition level nor the memory retention in trained bees. Appetitive olfactory learning performances in honey bees have several determinants, the main ones being the strength of the unconditioned stimulus (US) and the salience of the conditioned stimulus. Both factors were carefully controlled in our conditioning protocols. Yet, NPAAs might have induced changes in appetitive motivation and/or in the ability to sense and process odours.

To date, whether bees can perceive SMs in nectars remains controversial. Several nectar SMs are commonly considered as unpalatable or bitter, despite the lack of clear evidence supporting that bees perceive them as such [45, 47]. Behavioural studies have reported contrasting results so far. Quinine, a common nectar alkaloid, is widely used as aversive US in classical differential conditioning protocols, but its aversive value depends on the conditioning context [52]. Harnessed honey bees do not avoid food containing SMs even at concentrations high enough to harm or kill them [21, 53]. Electrophysiological studies failed to identify gustatory receptors firing in response to compounds bitter to human taste, but inhibition of sugar-sensing neurons has been reported in some cases [47]. Bees may thus indirectly perceive floral nectars as less sweet, rather than directly detecting SMs. However, this possibility fits poorly with our results, given that both GABA and β-alanine improved rather than weakened the acquisition level of bees encountering these NPAAs in the reinforcement offered during training. Improved acquisition in appetitive learning also did not reflect a generalized enhancement of feeding because in this and other studies [36] caged bees did not consume more food enriched with NPAAs. Moreover, in one assay freely moving bees did not prefer NPAA-free sucrose solutions. Overall, these results strengthen the view that honey bees, similarly to other generalist pollinators, have a low sensitivity for plant SMs. They also strongly suggest that the enhancement in learning observed in this study was not due to gustatory preferences for GABA or β-alanine.

The modulation of olfactory memory formation by NPAAs could be explained by a modified ability to sense and process odours. GABA is the most prominent inhibitory neurotransmitter in the honey bee primary olfactory brain centres, the antennal lobes (ALs) [32], and pharmacological application of GABA to the ALs abolishes the odour-evoked responses of projection neurons (PNs) [33]. We did not analyse the impact of NPAAs on the odour representations in the ALs, but we suggest that explanations via NPAA-induced changes of CS sensing can be cautiously excluded as our conditioning protocol used highly concentrated pure odorants. However, *calcium* imaging assays [54] will be crucial to rule out the hypothesis that NPAAs ingestion alters odour response patterns in the ALs.

Nectar NPAAs may enhance bees’ learning performance by pharmacologically affecting their central nervous system. Other nectar SMs directly interfere with bees’ neural activity and alter their behaviour [19, 20, 26]. For instance, caffeine strongly enhances long-term memory by modulating cholinergic neurons’ activity in the brain, which facilitates the association between the CS and the sucrose reward [19]. GABA, β-alanine, and taurine are important neuromodulators in the insect brain, and they may interact with bees’ neural activity soon after ingestion [30, 32, 55]. GABA, in particular, is the most abundant inhibitory neurotransmitter in the bee brain [56]. GABAergic neurons are located throughout bees’ central nervous system and peripherally at the neuromuscular junction [56, 57]. In the ALs, GABA signalling modulates the input received from antennal chemosensilla and the output sent to higher order brain structures [33, 58]. GABAergic feedback networks also regulate the firing response of Kenyon cells in the mushroom bodies (MBs) [59, 60], which are primary centres for multimodal integration and critical areas for learning and memory [61]. GABA modulates both ionotropic (GABA_A_) and metabotropic (GABA_B_) receptors. GABA_A_ receptors can also be pharmacologically activated by β-alanine and taurine [55, 62], a fact that might explain the similar effects induced by these NPAAs. As the effect on learning occurs within minutes of ingestion, we argue that these NPAAs may rapidly reach the haemolymph and the brain, and thus be able to directly interact with different neural networks by binding to receptor proteins of specific neurons.

NPAA ingestion two hours prior to conditioning caused impairment in acquisition, yet GABA did not affect mid-term memory retention, and taurine and β-alanine enhanced it. These results are puzzling because acquisition and memory retention performance are typically correlated. The fact that some NPAAs affected acquisition while leaving specific memory recall intact implies that bees acquired the odour-reward association during training even though they did not show appetitive responses. Differential effects on learning and memory have also been observed for acute doses of caffeine prior to or during training [51]. A potential explanation for our results is that these NPAAs temporarily decreased appetitive motivation in bees, most likely by inducing short-lasting satiation which then vanished at the time of the memory retention test. This hypothesis is partially supported by the fact that bees fed GABA exhibited significantly less appetitive responses during both conditioning and the memory test. Moreover, in our feeding assay bees consumed less taurine-laced solution over a prolonged period of time. NPAAs are by definition devoid of any nutritional value, but whether they induce satiation in insects is currently unknown. The epithelial endocrine cells of the midgut respond to many ingested substances, including toxins, by secreting several neuropeptides and hormones that finely regulate feeding behaviours [63]. It is worth noting that β-alanine and taurine selectively enhanced the specific memory and not the general response (NOd) of the bees. This evidence forces us to hypothesize that, in addition to changes in hunger state, other processes may underpin NPAAs’ effects on memory retention. As described above, these compounds have the potential to directly interact with a bee’s central nervous system shortly after ingestion. Future studies should aim at further exploring the complex mechanisms of action of nectar NPAAs on bees’ behaviour and cognition through pharmacological, genetic and neuroimaging techniques.

Our study is the first reporting that another class of SMs, nectar non-protein amino acids, interacts with learning and memory in a generalist pollinator. This is not surprising when we consider that NPAAs are one of the most common classes of SMs in floral nectars, that they are most frequently found in hymenopteran-pollinated plant species, and that at least some of them directly bind to neurotransmitter receptors on the membrane of insect neurons [5, 11]. Yet, so far, little work has been done to better understand their ecological role and their physiological mode of action. Several nectar SMs have been shown to increase plant fitness by increasing pollinators’ visitation rate or by filtering out undesirable pollinators [14, 16, 64]. Our results suggest that also NPAAs in floral nectar may increase plants’ reproductive success by facilitating learning of relevant flower features by pollinators. Therefore, we suggest that nectar NPAAs may be a cooperative strategy adopted by plants to attract beneficial pollinator, while favouring nectar transfer among conspecific flowers. Future studies should investigate whether these substances in nectar lead to suboptimal foraging choices by pollinators, in terms of reward value or floral constancy, as observed for other nectar secondary metabolites [20]. Research on plant-pollinator interactions is revealing increasingly complex ecological patterns involving multiple components. Our study highlights the need to study the role of other often overlooked classes of SMs in the regulation of pollination services received by plants. Future work should validate our results in ecologically relevant scenarios, as well as extend the study to other classes of SMs, alone and in combination, and to other species of pollinators.

## Material and Methods

### Artificial nectars

We tested five nectar non-protein amino acids: γ-amino butyric acid (GABA), β-alanine, taurine, citrulline and ornithine (Sigma-Aldrich, Italy). For each NPAA, we used a concentration reported to be within the upper limit of its natural range (GABA, 734 nmol ml^−1^; β-alanine, 267 nmol ml^−1^; citrulline, 283 nmol ml^−1^; taurine, 324 nmol ml^−1^ and ornithine, 323 nmol ml^−1^) [11]. Sucrose diluted in deionized water (Milli-Q system, Millipore, Bedford, USA) was used to prepare sucrose solutions. A 30% sucrose solution (w/w) was used as control. An identical 30% sucrose solution containing one of the five NPAAs was used as experimental solution.

### General methods (harnessing/housing)

Honey bee foragers (*Apis mellifera ligustica*) were collected while landing on a feeder providing 50% sucrose solution (w/w) and placed about 10 meters apart from an apiary consisting of six colonies (Department of Biology, University of Florence, Italy). For learning experiments (Exp 1, Exp 4 and Exp 5, see below), bees were immediately brought to the laboratory, cold anaesthetized, and individually harnessed in 3D-printed tubes using a small strip of duct tape placed in between the head and the thorax. A small drop of low-temperature melting wax was placed behind the head so that bees could freely move only their antennae and mouthparts [42]. After recovery, bees were fed with 5 μl of 30% sucrose solution to equalize the level of hunger and kept resting for two hours in a dark and humid place at room temperature (24 ± 2°C). For the feeding preference assay (Exp 2) and the mortality assay (Exp 3), bees were collected at a feeder as described above and immediately housed in customized 15 ml centrifuge tubes and in 250 ml Plexiglass cages respectively (see below for detailed information).

### Exp 1 – Chemo-tactile conditioning of the proboscis extension response (PER)

The appetitive response of the proboscis extension (PER) can be conditioned by repeatedly pairing a neutral stimulus with an unconditioned stimulus (US) that innately elicits the response so that, at the end of the training, the neutral stimulus will be conditioned (CS) and will evoke the response by itself [42, 43]. To test whether bees could detect natural concentrations of nectar NPAAs through their antennae, we used a chemo-tactile differential conditioning procedure [48]. Bees were trained to discriminate a chemo-tactile rewarded stimulation (CS+) from a chemo-tactile punished stimulation (CS-) over 10 trials (five CS+ trials and five CS-trials). Stimuli were presented in a pseudorandom sequence with a 12-min inter-trial interval. We used sucrose solution (30%, w/w) and NaCl solution (3 M) delivered by a toothpick to the bees’ antennae and proboscis as appetitive and aversive unconditioned stimulus (US) respectively [52]. Neutral/conditioned stimuli were prepared immediately prior to the experiment and consisted of 5 μl of either water or NPAA-laced water pipetted on small filter papers, presented to the bees’ antennae using a micromanipulator (WPI, MM-33). Each acquisition trial lasted 30 s. It consisted of a 14 s familiarization phase with the experimental context, a 6 s forward-paired presentation of the CS (either water or NPAA) and the US (either sucrose or salt respectively), (CS and US presentations lasted 6 s and 3 s, respectively, with a 3 s overlap) and a 10 s resting phase in the setup. For each bee, we noted PER occurrence during CS-only phase over the 10 conditioning trials. Prior to the experiment, all bees were allowed to drink water *ad libitum* so as to rule out the effects of thirst on appetitive responses [65]. An initial antennal stimulation with 30% sucrose solution was also performed to verify that the bees could properly extend their proboscis. Bees that failed to respond to this initial stimulation were discarded from the experiment. A total of 377 bees were successfully tested (GABA (n = 76), β-alanine (n = 81), taurine (n = 69), citrulline (n = 79) and ornithine (n = 72)).

### Exp 2 – Taste aversion/preference assay

In a second experiment, we investigated whether nectar NPAAs influenced feeding behaviour of bee foragers, adapting a protocol described elsewhere [49]. Bees were collected at the feeder using customized 15 ml centrifuge tubes, immediately fed with 5 μl of 30% sucrose solution and kept resting for one hour in a dark and humid place at room temperature (24 ± 2°C). Each customized tube contained a steel mesh that allowed the bee to freely walk. The tube was closed by a 3D-printed cap equipped with two 4 mm holes designed for hosting two 100 μl glass capillaries delivering two solutions: a 30% sucrose solution and an identical solution containing one of the NPAAs. The position of the microcapillaries (i.e. left *vs.* right) was balanced between bees to avoid any directional bias. The capillaries were connected to a 1 ml syringe by a silicon tube (6 mm length, 4 mm inner diameter), which allowed constant maintenance of the solutions at the tip of the microcapillary while bees were feeding. A micromanipulator kept the capillaries in the right position. After one hour of rest, the bees inside the tubes were placed into a polystyrene holder to screen off any visual stimuli and left to familiarize themselves with the apparatus for 5 minutes. Prior to the test, a small droplet of each solution (2.5 μl of either 30% plain sucrose or NPAA-laced 30% sucrose solution) was pipetted on the tip of the respective capillary and bees were allowed to explore and feed on the droplets. Bees that did not consume both droplets within five minutes were discarded from the experiment. Once the bees consumed the two droplets and moved away from the capillaries, they were immediately filled with the test solutions and put back in position. Bees’ activity was monitored for two minutes after the first contact with one of the two solutions. At the end of the test we took a picture of the two microcapillaries and measured the amount of solution consumed using the image processing software ImageJ (version 1.48). A total of 272 bees were successfully tested (GABA (n = 56), β-alanine (n = 46), taurine (n = 55), citrulline (n = 57) and ornithine (n = 58)).

### Exp 3 – Influence of NPAAs on feeding and mortality

A total of 900 bees were collected, cold anaesthetized, and housed in 250 ml Plexiglass cages in groups of 10 individuals [50]. Each container was equipped with two 20 ml syringes providing food *ad libitum*. Syringes were deprived of the luer cone to provide bees with easy access to the food and simultaneously prevent any sugar solution leakage. For each NPAA, three experimental conditions with six replicates each were run in parallel: 1) two syringes providing plain sucrose solution (S-S); 2) two syringes providing NPAA-laced sucrose solution (NPAA-NPAA); 3) one syringe providing plain sucrose solution and another one providing NPAA-laced sucrose solution (S-NPAA). To control for solution evaporation, an additional empty cage equipped with two 20 ml syringes was also maintained under identical conditions over the course of the experiment. Dead bees were counted and syringes were weighed daily to quantify mortality and sucrose consumption, respectively. Daily sucrose consumption was normalized as a function of the number of living bees in each container on a given day. The experiment lasted until all bees died (up to 10 days).

### Exp 4 – Contextual absolute olfactory learning

To test whether nectar NPAAs modulate bees’ ability to learn and memorize floral cues, we used an absolute olfactory conditioning procedure [42, 43]. Training consisted of four pairings of a neutral odorant (CS) with 30% NPAA-containing sucrose solution (experimental group) or 30% plain sucrose solution (control group). For each condition, we trained two groups in parallel using two different odorants as CS to avoid odour-specific biases in learning and to test specific memory after conditioning [42]. We chose 1-Hexanol and Nonanal as odorants since they are easily distinguishable by honey bees [66]. Prior to the test, bees were tested for intact PER by stimulating their antennae with 30% sucrose solution. Bees that did not respond to this initial stimulation were immediately discarded from the experiment. Odorants were provided to the bees by an automated odour releaser controlled by the microcontroller board Arduino Uno, which provided a constant airflow of clean air and allowed an efficient temporal control of the odour stimulation. An exhaust system ensured that odorants were removed from the apparatus through a hole placed behind the subject. Each acquisition trial lasted 30 s. The trials consisted of a 12 s familiarization phase with the automated odour releaser and the experimental context, a 6 s forward-paired presentation of the CS and the US (odorant and sucrose presentations lasted 4 s and 3 s, respectively, with a 1 s overlap) and a 12 s resting phase in the setup. A 12 min inter-trial interval was used. For each NPAA, we also ran in parallel an *explicitly unpaired* group to determine whether the increase in conditioned responses in the experimental and control groups were due to true associative learning [42, 43]. In this case, bees experienced four CS (either 1-Hexanol or Nonanal) and four US (either NPAA-containing or plain sucrose solution) presentations as the other groups, but in the absence of pairing: the US was provided 10 seconds before CS onset. Thus, bees in this group could not establish any associative link between CS and US. For each individual, we noted PER occurrence during the CS-only phase over the four conditioning trials. After conditioning, bees were kept resting for two hours in a dark and humid place before the memory retention test. Memory retention was assessed by presenting two odours to the trained bees: the conditioned odour (CS, either 1-Hexanol or Nonanal) and a novel odour (NOd, either Nonanal or 1-Hexanol, depending on CS). Odours were presented with an inter-trial interval of 12 min and the order of presentation of the CS and NOd was randomized between bees. Using the NOd allowed for testing of CS-specific memory. For each bee, we calculated specific memory as the difference between the response to the CS and the NOd during the memory retention test. A total of 867 bees were successfully trained (GABA (*paired group*: control, n = 46, exp, n = 52; *unpaired group:* control, n = 38, exp, n = 41), β-alanine (*paired group*: control, n = 62, exp, n = 66; *unpaired group:* control, n = 41, exp, n = 42), taurine (*paired group*: control, n = 38, exp, n = 38; *unpaired group:* control, n = 40, exp, n = 39), citrulline (*paired group*: control, n = 41, exp, n = 44; *unpaired group:* control, n = 41, exp, n = 43) and ornithine (*paired group*: control, n = 37, exp, n = 39; *unpaired group:* control, n = 39, exp, n = 40)).

### Exp 5 – Post-feeding absolute olfactory learning

NPAAs might also have post-ingestive effects that regulate hunger signals, eventually promoting or reducing bees’ appetitive motivation and, in turn, modulating associative learning [67]. To test this hypothesis, we ran a series of additional experiments in which bees were fed with either 5 μl of NPAA-laced sucrose solution or 5 μl of plain sucrose solution two hours prior to conditioning. After resting, bees were subjected to an absolute olfactory conditioning procedure as described above. In this case, we used 30% plain sucrose solution as reinforcement (US) and either 1-Hexanol and Nonanal as CS. An *explicitly unpaired* group was also performed to verify for true associative learning [42, 43]. After conditioning, bees were kept resting in a dark and humid place and tested two hours later for memory retention. The memory test was identical to the one described in Exp 4. We calculated specific memory as the difference between responses to the conditioned odorant (CS) and the novel odorant (NOd) during the memory test for each bee. A total of 737 bees were successfully trained (GABA (*paired group*: control, n = 29, exp, n = 35; *unpaired group:* control, n = 32, exp, n = 34), β-alanine (*paired group*: control, n = 38, exp, n = 35; *unpaired group:* control, n = 39, exp, n = 32), taurine (*paired group*: control, n = 37, exp, n = 36; *unpaired group:* control, n = 37, exp, n = 36), citrulline (*paired group*: control, n = 38, exp, n = 43; *unpaired group:* control, n = 40, exp, n = 42) and ornithine (*paired group*: control, n = 40, exp, n = 40; *unpaired group:* control, n = 37, exp, n = 37)).

### Statistical analysis

In both the tactile learning (Exp 1) and the associative olfactory learning assays (Exp 4 and Exp 5), responses to CS (PER: 1/0) of individual bees were analysed using repeated-measure ANOVA. In all cases, independent models were used for each NPAA. We ran a series of generalized linear mixed models (GLMMs) with a binomial error structure - logit-link function, *glmer* function of R package *lme4*. When necessary, models where optimized with the iterative algorithm *BOBYQA* or *Nealder-Mead*. In the models run for the tactile learning assay (Exp 1) “*bee response*” was entered as dependent variable, ‘*CS*’ as fixed factor and “*Stimulation trial*” as a covariate. In the models for the acquisition phase of the associative olfactory learning assays (Exp 4 and Exp 5), ‘*bee response*’ was entered as dependent variable, ‘*treatment*’ *(NPAA-laced sucrose solution vs plain sucrose solution)* and ‘*CS*’ *(1-Hexanol vs Nonanal)* as fixed factors, and ‘*conditioning trial*’ was used as covariate. In all models the individual identity “*IDs*” was entered as a random factor to account for repeated measures. Interactions were evaluated in all full models and reported when significant. In all models, we retained the significant model with the highest explanatory power (i.e., the lowest AIC value). A series of χ^2^ tests were used to compare the responses to the CS, the NOd and the specific memory performances of bees rewarded (Exp 4) or pre-fed (Exp 5) with plain sucrose and that of bees rewarded/pre-fed with the NPAAs. In the feeding preference assay (Exp 2), we calculated a preference index for NPAAs by subtracting the residual volume of the liquid inside the control capillary from the one inside the NPAA capillary measured at the end of test. A positive value of the index indicates a preference for NPAAs while a negative value indicates a preference for plain sucrose solution. For each NPAA, we ran a one-sample Wilcoxon test to compare the observed preference index against the hypothetical value of 0 which indicates a lack of preference. In the feeding and mortality assay (Exp 3), sucrose consumption was evaluated at 24 h and at the end of the experiments using Mann-Whitney U test. For the statistical evaluations in the survival experiments, we ran the Breslow Statistic (Mantel–Cox Test) using the *Cox* function of R package *Survival*. In the regression model “*diet*” (Sucrose-Sucrose, Sucrose-NPAA and NPAA-NPAA) was entered as fixed factor while “*cage identity*” as random factor. All analyses were performed with R 3.4.2.

## Acknowledgments

Thanks are due to Lara Sassoli for help with experiments, to Zoe Korzy Wild for help in proofreading the manuscript and to Ken Cheng for the useful comments on the manuscript.

## Funding

This work was supported by the Italian Ministry of Education, Universities and Research (MIUR) (Rita Levi Montalcini Fellowship for Young Researchers provided to D.B. Project ID PGR15YTFW9) and the University of Florence.

## Author Contributions

D.B. conceived the study and designed the experiments. E.P., S.S. and G.B. performed most of the experiments. D.B. carried out data analyses and prepared figures. D.C. wrote the first draft of the manuscript. D.B. and D.C. wrote the final version of the manuscript. D.B. obtained the funding for the research.

## Competing interest

Authors declare no competing interests.

## Notes

### Competing Interest Statement

The authors have declared no competing interest.

